# Structure of the type VI secretion protein VgrS from *Salmonella* Typhimurium

**DOI:** 10.1101/2024.12.20.629533

**Authors:** Kartik Sachar, Matthew Van Schepdael, Karsen L. Winters, Gerd Prehna

## Abstract

Enteric bacterial pathogens employ various strategies to colonize the intestine and cause diseases ranging from gastroenteritis to systemic infections. For example, *Salmonella enterica* utilizes a nanomachine known as the type VI secretion system (T6SS) to facilitate colonization of the host gut. However, the varied mechanistic details of how the T6SS is loaded with effector proteins remains to be elucidated. Here, we present an X-ray crystal structure of the *Salmonella* Typhimurium VgrG (VgrS) that serves as platform for T6SS effector loading. Compared to other known structures of VgrG proteins, the VgrS trimer adopts an alternative open conformation composed of a domain-swap between the monomers in the gp27 region. Additionally, a comparative structural analysis of VgrS with other VgrG proteins reveals molecular variations that may contribute to specific effector loading mechanisms. Our structural data and molecular analysis highlight the observation that the T6SS of each bacterial species or strain is unique.

## INTRODUCTION

Bacterial antagonism and the competition for essential nutrients and space is a primary factor that shapes microbial community structure and dynamics. Consequently, bacteria have developed sophisticated multi-protein nanomachinery to engage in interspecies competition and eliminate rival bacteria. *Salmonella enterica*, a major cause of human gastroenteritis, employs such a competition mechanism named the type VI secretion system (T6SS) ^1–3^. The T6SS is a dynamic nanomachine that Gram-negative bacteria use to inject toxic effectors to kill rival bacteria in a contact-dependent manner ^4^. *S. enterica* encodes several T6SS across different *Salmonella* Pathogenicity Islands (SPIs) and serovars ^3,5^. However, each *S. enterica* subtype and serovar has unique T6SS molecular adaptations that result in different clinical manifestations and host specificity ^3,5^.

The current T6SS model includes 13 core proteins (TssA-M, *Pseudomonas* nomenclature), divided into three components: a membrane-anchoring complex, a phage tail-like structure, and a baseplate complex ^6^. The membrane-anchoring complex secures the apparatus to the bacterial membranes, while the baseplate acts as a scaffold for assembling the tail-like structure ^7–11^. The phage tail-like structure functions as the site for effector recruitment and consists of four proteins: TssB, TssC, Hcp, and VgrG ^12,13^. The TssBC proteins form a sheath that assembles around the homo-hexameric Hcp tube. Upon full extension of the tail, the TssBC sheath contracts ^14^. This propels the Hcp tube, VgrG, and their bound effectors into an adjacent cell ^13^.

The primary function of the T6SS is to secrete protein effectors, which exert diverse toxic effects on the targeted prey cell ^15^. A critical step before secretion is the assembly and loading of effectors onto the T6SS apparatus. There is no singular structural feature that defines T6SS effector loading, with effectors often associating with the protein VgrG that forms the T6SS puncturing tip ^16–21^. VgrG proteins can be covalently linked to an effector, or effectors can be non-covalently loaded onto VgrG by the use of adaptors and chaperones ^16,22–25^. For example, some effectors require cone-shaped adaptor domains termed PAAR, ‘PAAR-like’ or PIPY, which bind the top of the VgrG β-prism and are said to ‘sharpen’ the T6SS ^26–29^. PAAR domains can also require a chaperone for loading, such as Eags (DUF1795) that load membrane protein effectors onto VgrG ^23^. Furthermore, different VgrG proteins can have unique structural modifications that serve to recruit adaptors. Namely, adaptors such as Tap(DUF4123) recognize a helix-turn-helix extension in their cognate VgrG to load diverse effectors ^30^. Additionally, VgrG can be extended by adaptor domains such as transthyretin-like (TTR) to recognize effectors ^16,31^. Regardless of mechanism, what is apparent is that VgrG serves as universal but modular platform for effector loading.

Given the extensive variations in effector loading, detailed molecular data is required to understand how proteins associate with the T6SS apparatus. As VgrG is a modular central platform, each VgrG provides insight into molecular adaptions for species-specific effector secretion. In the case of *Salmonella* Typhimurium, it only encodes one VgrG within the T6SS found in SPI-6, but has a diverse set of several effectors that must be loaded for secretion ^1,32,33^. As such, the mechanism by which multiple different effectors are loaded onto the Salmonella T6SS for secretion remains unclear.

To help address the questions of effector loading, we have solved an X-ray crystal structure of the *Salmonella* Typhimurium VgrG named VgrS (STM0289) ^1,3,34^. The VgrS crystal structure reported here shows an alternative open conformation relative to other VgrG proteins and reveals molecular features likely unique to *S.* Typhimurium T6SS effector loading. Our data further highlights that each T6SS is unique and specifically modified for the needs of each bacterial species.

## RESULTS

### Genetic Context of VgrS in *Salmonella spp*

VgrS is encoded within Salmonella Pathogenicity Island-6 (SPI-6), which houses all core T6SS genes ^1,3,5,34^, and four known effector/immunity pairs: *tae4/taiA, tlde1a/tldi1a*, *rhs1/rhsI1*, and *rhs2/sciX* (Figure 1A) ^1,2,33,35^. Among these effectors, only the PAAR-containing Rhs1 has a defined mechanism for loading onto VgrS ^23,36^. Furthermore, when compared to other *Salmonella* serovar T6SSs the genetic organization the SPI-6 T6SS shows variability, especially among the repertoire of effectors (Figure 1B) ^3^. The VgrS from *S.* Typhimurium and *S.* Typhi appear closely related but diverge in sequence from *S.* Gallinarum and *S.* Dublin, which contain a gp5 extension absent in both *S.* Typhimurium and *S.* Typi (Figure S1). Interestingly, *S.* Typhimurium and *S.* Typhi are major human pathogens whereas *S.* Gallinarium and *S.* Dublin appear adapted for chickens and cattle ^1,3,37,38^. Given this, we hypothesized that each VgrS must have unique molecular features to facilitate serovar-specific effector loading. As such, we pursued an experimental structure of VgrS to observe potential structural adaptations related to secretion.

**Figure 1:**
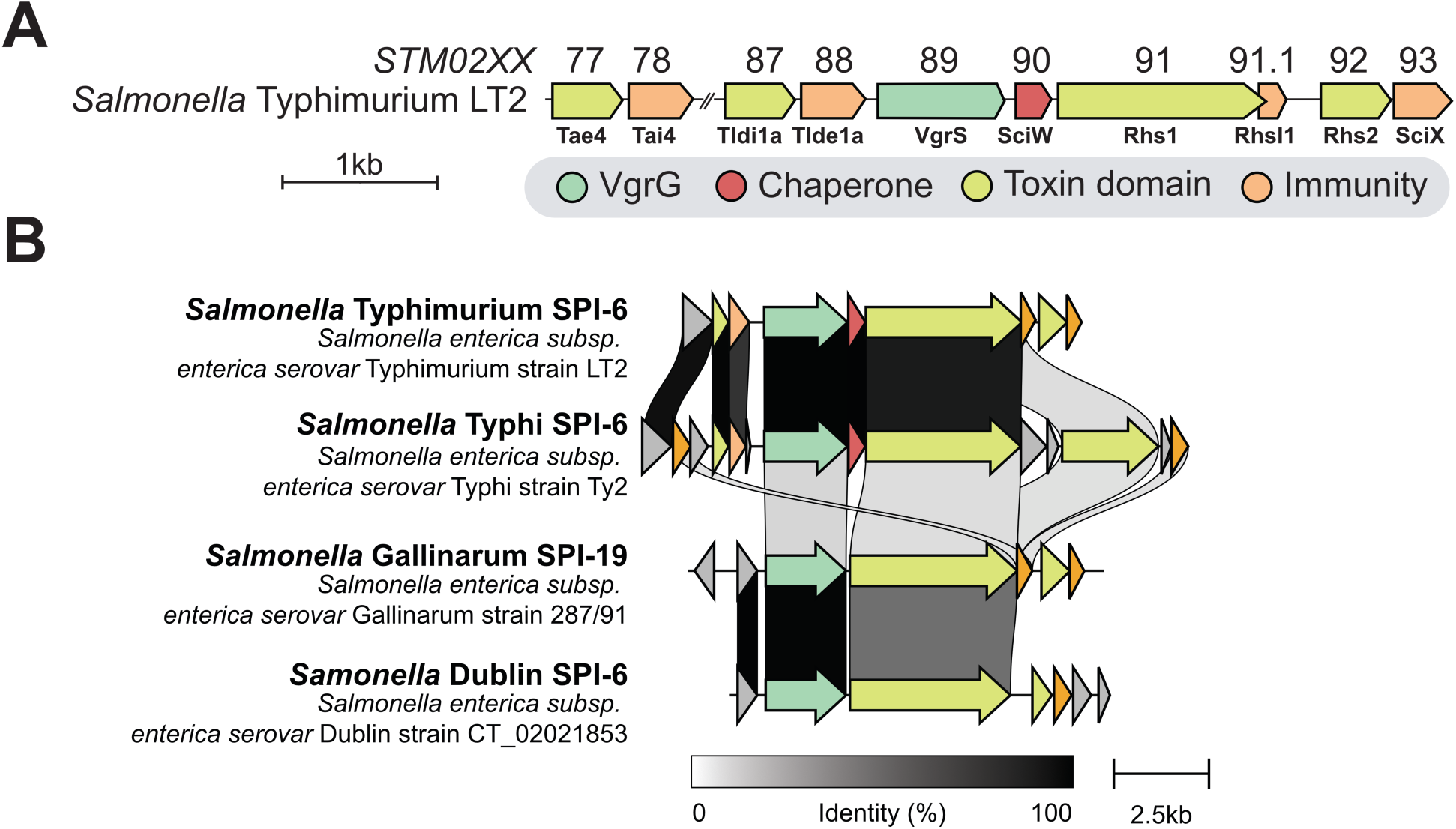
Schematic representation of *Salmonella* T6SS gene clusters. **A)** The genes surrounding VgrS (STM0289) in *S.* Typhimurium LT2. **B)** Schematic representation of *Salmonella* T6SS gene clusters. Alignment showing the sequencing identity of genes surrounding VgrS (STM0289) in four *Salmonella* serovars. VgrS is also known as VgrG.

### X-ray crystal structure of *S.* Typhimurium VgrS

Full-length VgrS produced insoluble material, but a truncated construct (residues 81-729) that removed a region of predicted disorder (Figure S1) was both soluble and purifiable ^39,40^. Diffracting crystals of VgrS (81-729) were obtained and the structure of VgrS was solved by molecular replacement to a resolution of 3.1Å using a previous solved model of VgrG from *Pseudomonas aeruginosa* (PDB: 6H3L) ^17^ (Figure 2 and Table 1). The data can be processed as P6_3_22 with a VgrS monomer as the asymmetric unit. However, the lower symmetry space group C222_1_ was chosen for refinement to have the biologically relevant VgrS trimer in the asymmetric unit (Figure S2A). The lower symmetry was also chosen as the interior of the trimer exhibited clear ligand density coordinated by each monomer (Figure S2B). In the crystal, the VgrS trimers pack tip-to-tip via their β-helix neck regions and the base of each trimer packs with the base (gp27 head) of a 2-fold rotated symmetry mate (Figure S2A). Strikingly, the resulting crystal packing features a rarely observed extremely high solvent content (82.22%), with a Mathews Coefficient of 6.91.

**Figure 2:**
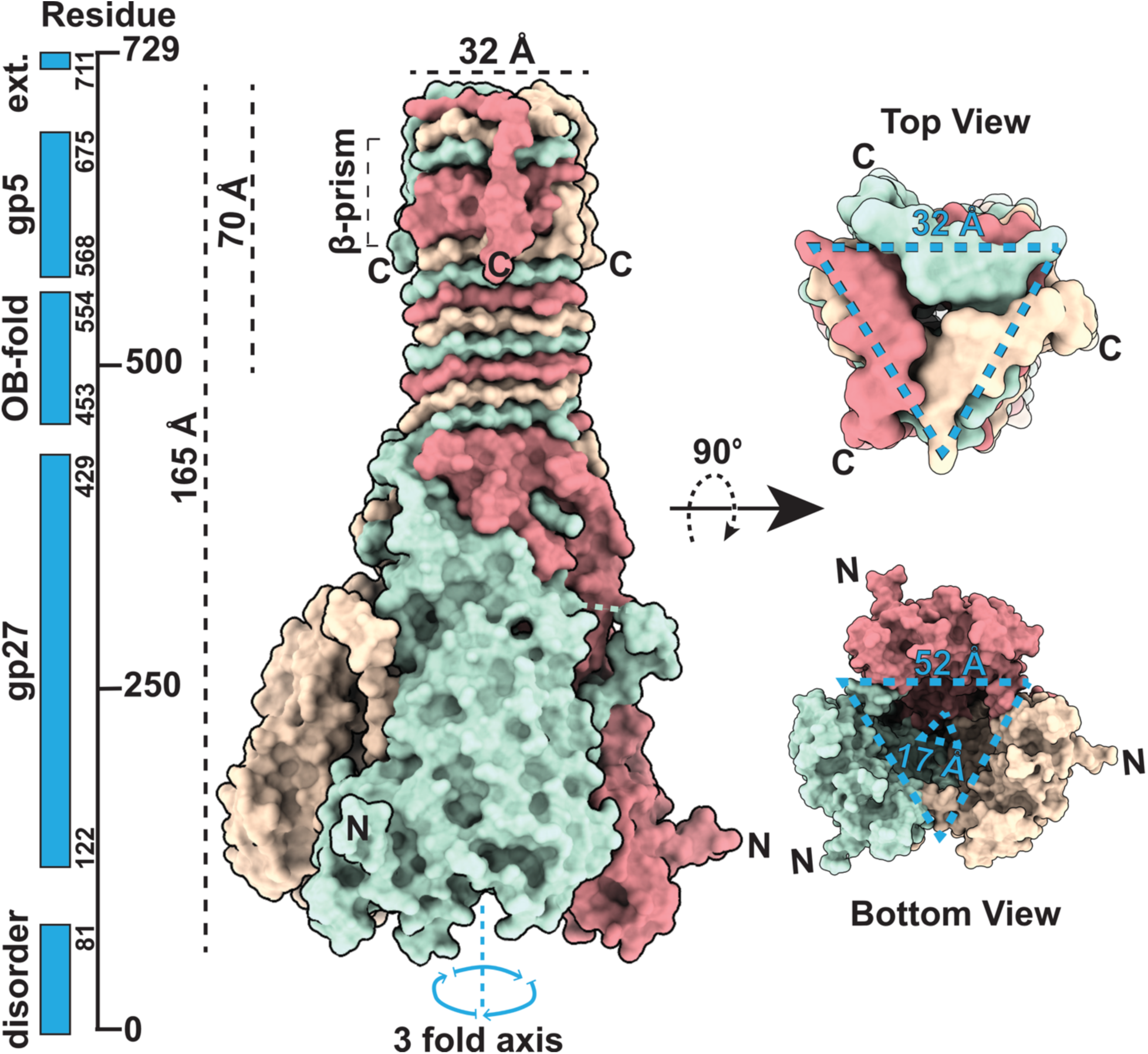
X-ray crystal structure of VgrS. Overall structural features of VgrS. Domain annotations for the VgrS sequence were generated using predictions from the CDD server and outlined by residue boundaries (left). The VgrS trimer is shown by surface representation with each monomer shown in a different color (middle). The top of the spike and base of the VgrS are shown and structural features measured (right).

**Table 1:**
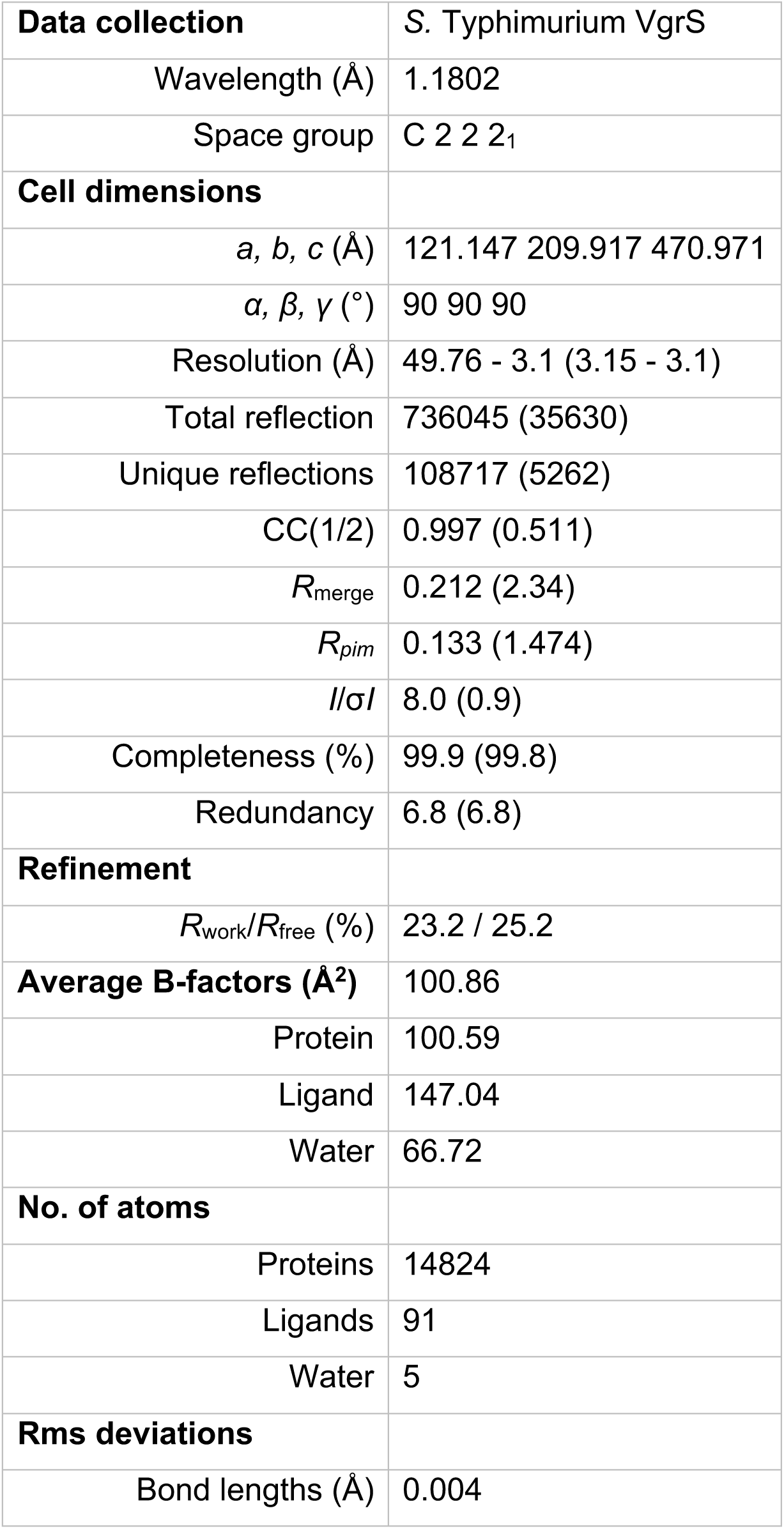
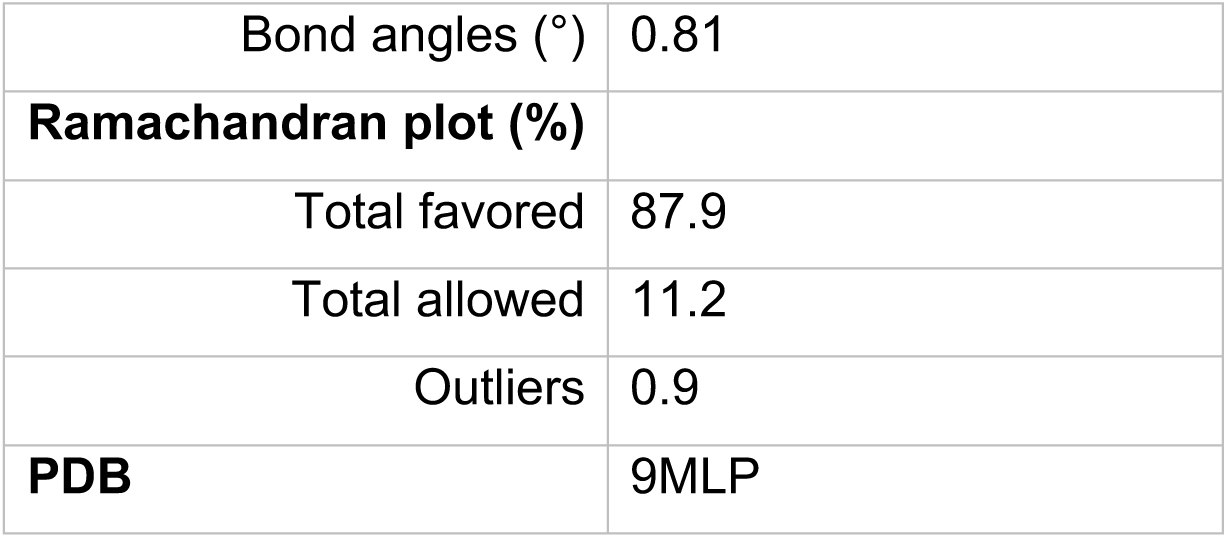
X-ray data collection and structure refinement for VgrS.

Similar to other VgrG proteins, VgrS is a trimer composed of domains that are structurally homologous to the T4 bacteriophage spike-protein ^41^. These domains include the head (gp27, residues 122 - 429), an OB-fold domain containing neck (residues 453-544), and the spike (gp5, residues 483 - 548) (Figure 2) ^16,20,21,36,41^. This includes a hollow base (gp27 head), and a spike made of a triple-stranded β-helix (alternating strands from each monomer) and an interleaved β-prism (Figure 2). The spike creates a flat triangular platform atop VgrS of ∼32 Å per side that serves as the binding site for PAAR domain containing effectors ^18,23,26,36^.

Despite these similarities, the structure of VgrS diverges from other known VgrGs. The first notable difference is that VgrS contains a domain swap between each monomer (Figure 3A and 3B). Additionally, the gp27 head domain cup-like structure of VgrS has a larger cavity than other VgrG proteins. The VgrS gp27 head measures approximately ∼52 Å at the base and tapering to about ∼17 Å near the top (triangular dimensions) (Figure 2 and Figure 3A, bottom views). Next, the VgrS platform contains a C-terminal tail extension (residues 711-729) that is approximately 31Å long and notably shorter than extensions found in other VgrGs (Figure 3A and Figure S1) ^16,17^. In particular, the molecular features of C-terminal extensions have been implicated in many T6SS effector loading mechanisms ^22,24,30,42,43^, suggesting a unique role (or lack of role) in VgrS effector interactions. Additionally, VgrS contains a domain swap where residues 319 to 366 form an elongated loop that contacts the subsequent monomer (Figure 2, Figure 3B, and Figure 3C top panel). This is in contrast to the known VgrG1 structures where the same region forms anti-parallel beta-sheets which pack within the same chain of the trimer (Figure 3A).

**Figure 3:**
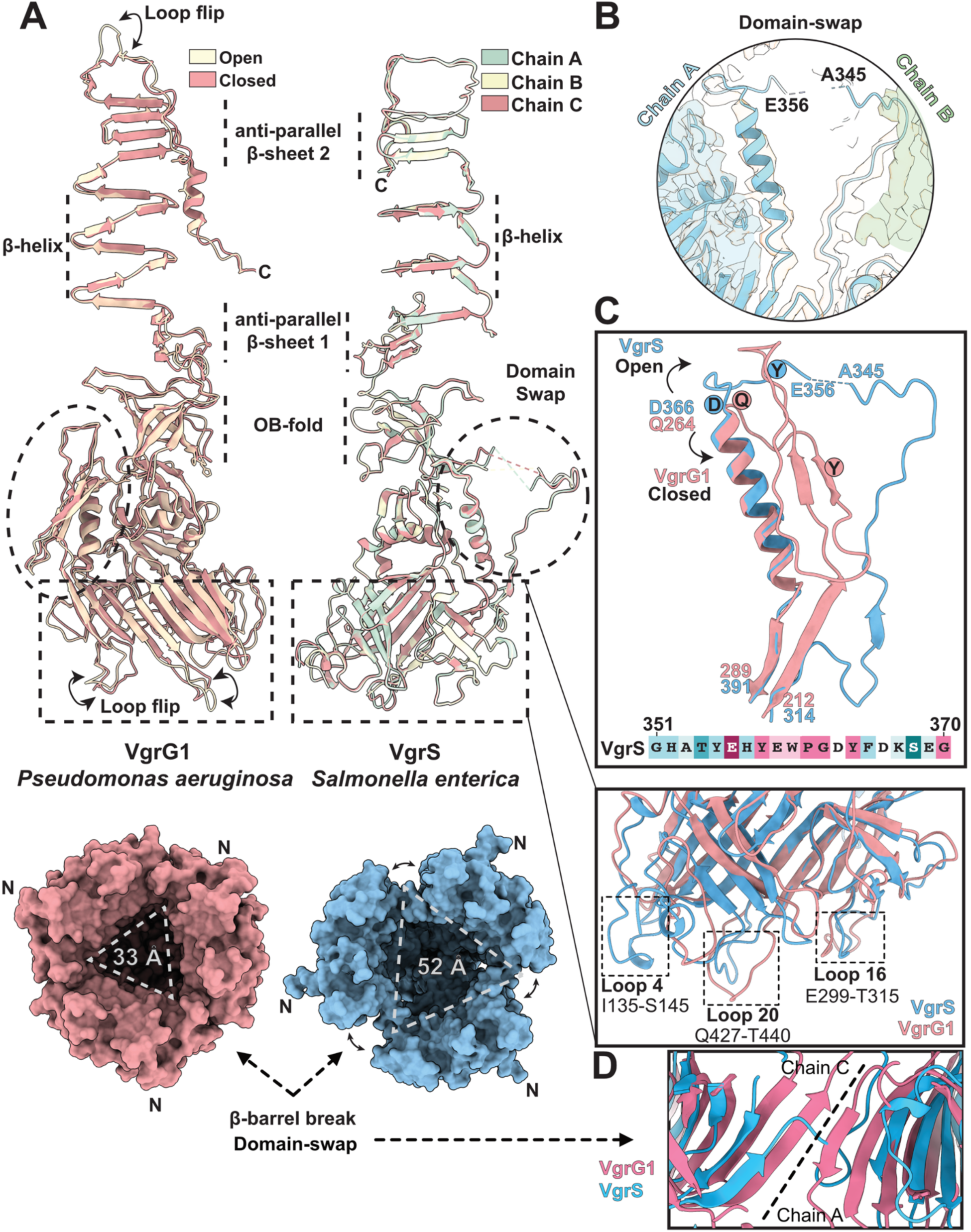
The VgrS domain swap expands the gp27 head base to adopt an open conformation. **A)** Comparative analysis of asymmetric units of VgrG1 and VgrS crystal structures. Top left: Overlayed chains of VgrS and VgrG1 in their “open” & “closed” conformation (PDB: 4UHV). Top right: Overlayed chains of each VgrS monomer. The size of the gp27 base is shown below each structure. **B)** Structure of the domain-swap region. A 2mFo-DFc density map illustrating the fit of the VgrS domain swap. **C)** Comparison of domain-swap region of VgrS with VgrG1 (PDB: 4UHV). VgrS is shown in blue and VgrG1 in pink. Conserved residues in the region are highlighted using ConSurf ^61^ and relative positions shown on the structures. Bottom: Differences in the loop region conformations important for Hcp binding. **D)** Break of continuous interchain beta-barrel in VgrS compared to VgrG1 (PDB: 6H3L) due to the domain swap. The interchain beta-barrel is formed by residues Q214-Q217 in chain C with C77 to V80 in Chain A of VgrG1.

During model building, we also identified electron densities at the β-helix and the interior of the gp27 cup that could not be modeled as water (Figure S2B). Given that VgrS crystallized in high concentrations of ammonium sulfate (1.5M), the exterior ligands could be explained by sulfate ions. However, the central density coordinated by all three monomers at residues R620 and K622 could not be initially identified. As such, we performed inductively coupled plasma mass-spectrometry (ICP-MS) to help determine the identity of the bound ligands. We used the Eag chaperone SciW as a negative control as it does not bind metals and the zinc-containing papain-like protease from SARS-CoV-2 (PLpro) as a positive control ^23,44,45^. ICP-MS results showed no significant enrichment of any tested metal ions in VgrS compared to the negative control (Figure S2C). Similar density was observed in VgrG1 (PDB: 4UHV) and VgrG1 (PDB: 4MTK) from *Pseudomonas aeruginosa* (Figure S2B) ^20^. The internal density for 4UHV was not modelled, but for 4MTK the density was modelled as sulfate in part due to the crystallization conditions (1.4M LiSO_4_). We therefore modeled this central density as a sulfate ion, though other anions like chloride remain possible (Figure S2D). Overall, coordination of an internal anion by each monomer of the β-helix could be a conserved feature of VgrG proteins. Additionally, a Fo-Fc difference map demonstrates the position with clear density for the domain swap showing that the built conformation is not due to modelling error or bias (Figure S2E).

### The VgrS head domain adopts an open conformation

A previous crystal structure of VgrG from *Pseudomonas aeruginosa* (PDB: 4UHV) shows that the head domain of VgrG1 can exist in two states, an “open” and “closed” conformation (Figure 3A) ^20^. In the open conformation, the cavity at the base of the head domain widens due to the movement of loops at the very base of VgrG. This in turn creates an uneven spike top, which is unfavourable for PAAR binding (Figure 3A, left). In contrast, the three chains of the VgrS trimer in our structure have identical conformations (r.m.s.d of 0.235 Å^2^) (Figure 3A, right). Comparing the VgrG1 open and closed conformations to VgrS, they align within the neck and spike (OB-fold and gp5) but differ in the gp27 base. This is especially pronounced at both the VgrS domain swap and in the conformations of the open/closed loops at the base (Figure 3B-C). VgrS differs from both the open and closed versions of VgrG1, with the comparison to the open VgrG1 highlighted. These differences also appear true for VgrS compared to all known VgrG structures (Figure S3A).

Instead of folding into anti-parallel β-strands as observed in other VgrG structures (Figure S3A), VgrS residues 319 to 366 (VgrG1 residues 220 to 264) form a loop that domain-swaps into the neighboring monomer. This makes all three monomers appear to ‘grasp’ the donated loop around the base of the trimer (Figure 2 and Figure 3B-C). Of note, VgrS residues A345 to E356 could not be modeled due to insufficient electron density likely indicating flexibility (Figure 3B and Figure S2E). We cannot rule out that the missing density is due to cleavage or degradation during crystallization. However, given that the conformation of the domain swap was the same in all three VgrS monomers, with the exact same portion of unmodelled residues, the observed conformation is unlikely to be due to non-specific cleavage or degradation. Furthermore, the domain swap results in the lack of a continuous β-barrel formed by all three monomers at the base of the gp27 fold that is characteristic of other VgrG proteins (Figure 3D). Overall, the domain swap between the monomers results in a broader base and introduces a solvent accessible gap between the monomers which in turn pushes VgrS into a conformation far wider than either the closed or open VgrG1 conformations (Figure 2 and Figure 3A). This domain-swap has only been observed in VgrS setting it apart from previously described VgrGs (Figure S3A) ^17,18,20,21^.

To further probe the potential conformational states of VgrS we generated an Alphafold3 (AF3) model of trimeric VgrS (VgrS_AF_) (Figure S3B). The predicted VgrS_AF_ model resembles the closed conformation of VgrG1 more closely than the crystal structure of VgrS (r.m.s.d. of 4.206 Å^2^ vs. 7.176 Å^2^). Specifically, the domain swap residues observed in the VgrS crystal structure are modelled to match the anti-parallel β-sheet of VgrG1 and other VgrGs. This includes the characteristic continuous β-sheet between all monomers within the gp27 fold. The base of the VgrS_AF_ model is also more compact than the crystal structure, resulting in a cavity of similar size to the closed conformation of VgrG1. (Figure 3A and Figure S3C). While this may represent training-set bias in AlphaFold3, it is possible that VgrS adopts both conformations represented by the crystal structure and the AlphaFold model. For VgrS, this conformational switch could be linked to conserved residues 356 to 366 (EHYEWPGDYFD) changing from an extended domain-swapped loop to a compact anti-parallel β-sheet similar to VgrG1 (Figure 3C, top panel).

We also examined the B-factors in the VgrG structures, which can reveal regions of flexibility ^46^. In VgrS, the highest B-factors were observed in the gp27 head domain at the domain-swap region and in the loop regions (Loop 4, 16, and 20) (Figure 3C, bottom panel and Figure S3D). This supports our hypothesis that VgrS may change conformation between a closed conformation as seen in VgrS_AF_, and the open form observed in our structure. Additionally, a comparison with a VgrG1 crystal structure shows that in the VgrG1 open conformation the domain swap region and loop regions also exhibit high B-factors (Figure S3D). Notably, the loop regions are thought to change conformation to interact with Hcp ^18,20^. However, this also suggests that the equivalent VgrS domain-swap region in all VgrGs may have the ability to change conformation.

### The molecular surface of VgrS reveals clues to effector loading

Given that VgrS diverges within *Salmonella* serovars (Figure 1 and Figure S1) and is structurally different from other VgrGs (Figures 2–3 and Figure S3A), we hypothesized that VgrS may have molecular features for *S.* Typhimurium specific effector loading. As such, we examined in detail the molecular surface properties of VgrS and its unique structural elements. These include the short C-terminal tail of VgrS, the uniquely large inner cup of the gp27 head domain, and the sulfate coordination ring near the OB-fold.

The molecular surface of VgrS exhibits limited conservation within the C-terminal tail and gp27 head domain but is highly conserved in the neck region (Figure 4A). Notably, the base of the gp27 head domain shows no conservation especially within the loop regions that interact with Hcp. However, the back of the hollow cup exhibits residue conservation and is interestingly a region of sulfate binding (Figure 4A and Figure S2B). Furthermore, this region becomes the highly conserved OB-fold motif followed by an anti-parallel β-sheet that is also a site of sulfate coordination. Specifically, conserved pairs of surface exposed electropositive residues Arg586/Lys597 and electronegative residues Glu601/609 bind sulfate (Figure 4A–B and Figure S2B). This conserved residue pattern in VgrS raises the possibility that these regions contribute to effector loading or interaction with an adaptor through electrostatic or salt-bridge dependent mechanisms. However, this observation remains speculative and would require further experimental investigation.

**Figure 4:**
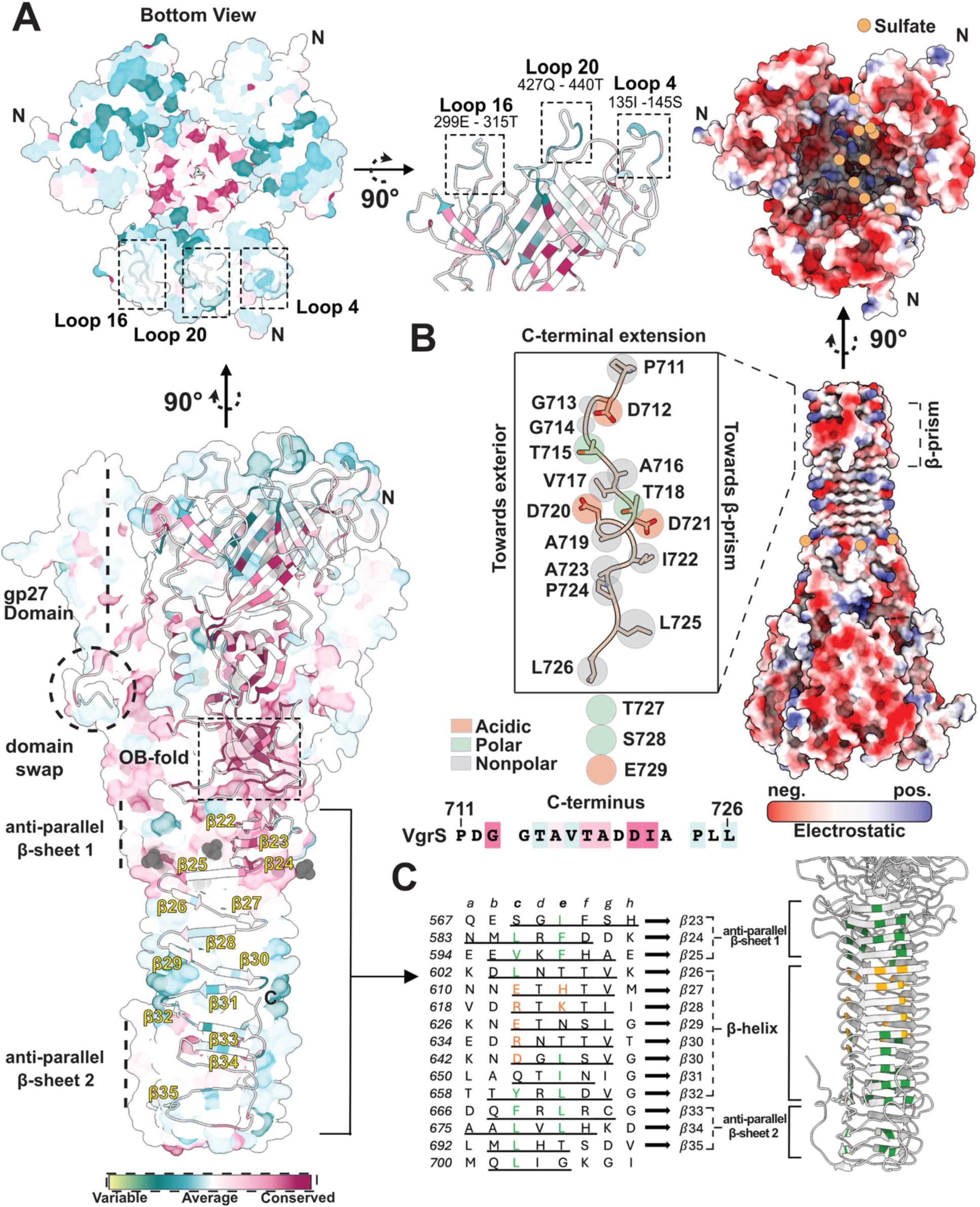
Molecular surface properties of VgrS. **A)** Surface conservation of VgrS as determined by the server ConSurf. Top: Molecular surface of the gp27 base with a rotated view of the Hcp binding loop regions showing they are unconserved. Bottom: Annotated molecular surface of VgrS. **B)** Relative electrostatic potential of the VgrS surface shown in red (negative) and blue (positive). A rotation showing the surface of the gp27 base is also shown with positions of sulfate ligands indicated (orange). The inset is a zoom in of the C-terminal extension highlighting acidic (red), polar (green), and nonpolar (grey) residues. The orientation of the residues is indicated as pointing either toward or away from the β-prism. Residue conservation of the C-terminal tail is shown underneath **C)** Manual sequence alignment of the spike β-strands in VgrS (underlined). Left: Residues are coloured orange and green as hydrophobic and hydrophilic, respectively. Right: The C-terminal spike region (white) coloured to show the repeat hydrophobic (green) hydrophilic (orange) hydrophobic (green) pattern.

In line with an electrostatic mediated effector loading model, we analyzed the electrostatic surface of VgrS to gain further insight. Overall, VgrS is highly electronegative with patches of electropositive residues that show residue conservation (Figure 4B).

Notably, there is a high concentration of electronegative residues on the external face and putative Hcp binding regions at the bottom of the gp27 head cup (Figure 4A-B). The internal face of the gp27 head cup also shows high electronegativity but exhibits an electropositive surface deep within the base. At the base of the cavity, multiple conserved His, Lys, and Arg residues (Figure 4A left) coordinate sulfate ions, suggesting a potential binding-partner surface yet to be determined (Figure 4B, rotated view top). In contrast, the uniquely short VgrS C-terminal extension contains several acidic residues. The C-terminal extension runs downward alongside the spike, with the loop folding to interact with the β-prism formed by the same chain so that residues D715, T718, D720, and P724 point away from the β-prism (Figure 4B, inset). It is possible that VgrS recruits *Salmonella* specific adaptors for effector loading through its unique electronegative extension given comparisons to other VgrGs ^22,30,42^.

Finally, we also examined VgrS for hydrophobic regions. Extensive hydrophobic interactions exist in the gp-27 head, the OB-fold neck domain, and the C-terminal spike. The spike region contains an internal repeating hydrophobic-polar-hydrophobic environment created by the side chains of residues pointing towards the centre of the β-helix (Figure 4C). The first anti-parallel β-sheet (β23-β26) shows mainly hydrophobic residues (green) followed by a β-helix (β27-β32) which contains hydrophilic inward-facing side chains. The remaining β-helix including the second antiparallel β-sheet (β33-β35) contains hydrophobic residues. This repeating pattern is characteristic of the spike region reported in VgrG1 in *P. aeruginosa* ^20^. However, overall VgrS does not have surface exposed hydrophobic patches further driving the hypothesis that effector loading is electrostatic mediated.

### Potential VgrS conformational changes in T6SS assembly

Current structural data of VgrG proteins bound to PAAR domains and/or Hcp report VgrG to also be in the closed conformation ^18,36^. This suggests that VgrS likely changes conformation (Figure 3A and Figure S3B) upon interaction with Hcp or potentially other T6SS proteins. For example, studies have shown that the flexible loops of Hcp interact with the dynamic loops at the base of a VgrG gp27 domain (Figure 3C bottom, Figure 4A) ^18^. Therefore, Hcp binding would likely require VgrS to close which would move the gp27 domain dynamic loops into position for interaction with the Hcp ring.

In *S.* Typhimurium, three copies of the *hcp* gene are present. These include *hcp1* (STM0276) and *hcp2* (STM0279) both located within SPI-6, and a distally located *hcp3* that is not present within SPI-6 ^47,48^. Figure 5A shows the previously solved crystal structure of Hcp2 (PDB: 5XEU) which exhibits a similar sized but smaller ring compared to other Hcp-fold structures (∼35 Å inner measurement) (PDB: 7YW0) ^18^. Additionally, we used AlphaFold3 to model both Hcp1 and Hcp2 as a hexamers, because Hcp1 (5XHH) is deposited as a monomer (Figure S4A). Note, the AlphaFold models closely resemble the Hcp2 crystal structure (r.m.s.d. 0.695 Å^2^) (Figure S4B). Both Hcps are also highly conserved with only 10 amino acid differences that are primarily located on the interior lumen of the hexameric rings. Given this, they exhibit extremely similar surface properties such as electrostatics and residue conservation of the flexible loops on their VgrS binding surface (Figure 5A). In line with this, when we used AlphaFold3 to dock Hcp1/2 with VgrS the model predicts that VgrS adopts a closed conformation so that the dynamic loops of the VgrS base might contact the flexible loops of the Hcp ring (Figure 5B and Figure S4C). Namely, when comparing the domain swap region, we see that AlphaFold chooses the closed form β-sheet for VgrS residues 356 to 366 and completes the gp27 base.

**Figure 5:**
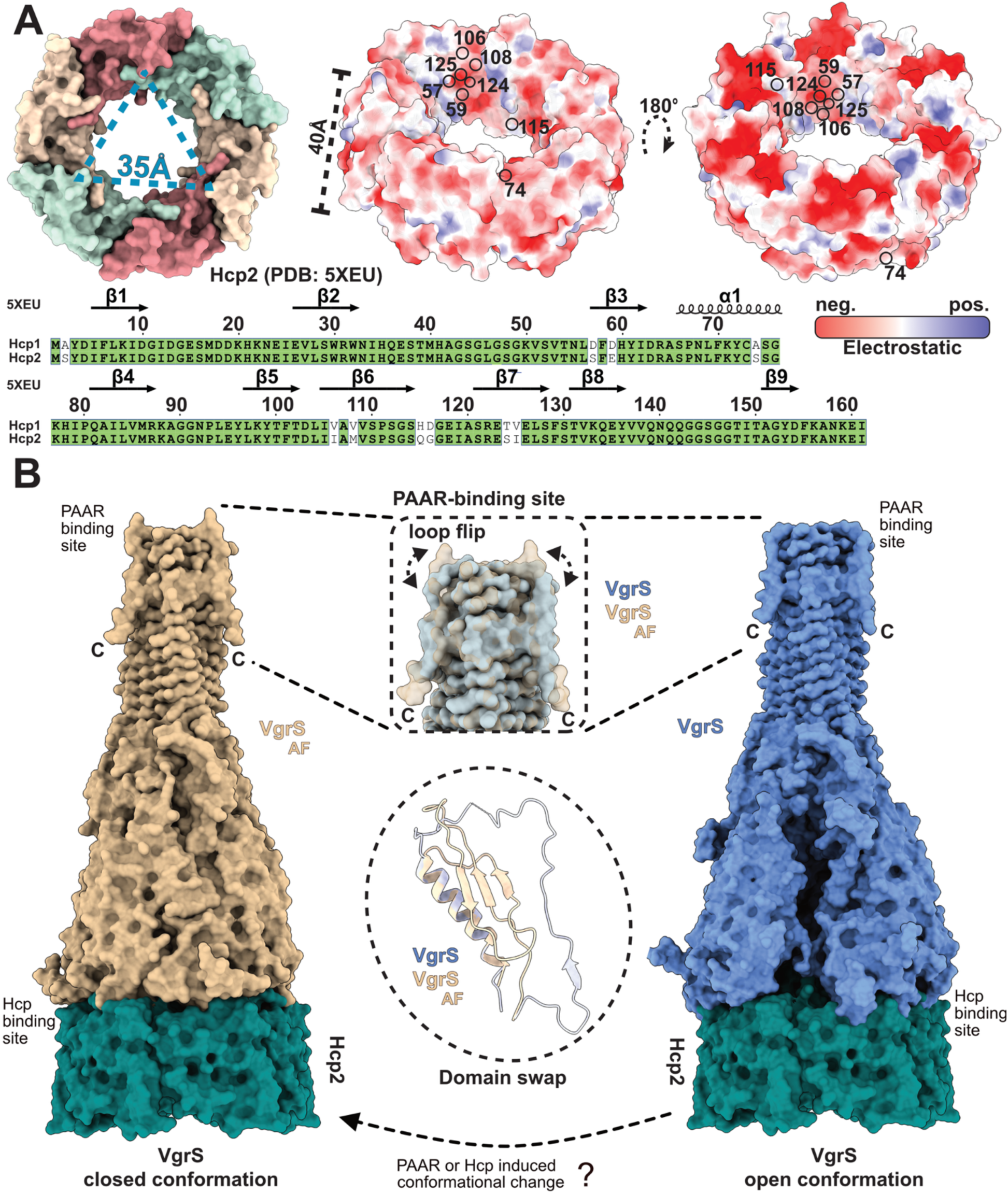
Comparison VgrS and a VgrS:Hcp model. **A)** The general hexameric structure of *Salmonella* Hcp proteins is shown on the left, with the triangular dimensions of Hcp2 (PDB: 5XEU) measured from the Tyr03-OH of every second chain. Middle and right show an electrostatic potential map of Hcp2 (PDB: 5XEU) and an AlphaFold hexamer of Hcp1, respectively. Bottom: Sequence alignment of Hcp1 and Hcp2, with the sequence differences highlighted on the electrostatic surfaces. **B)** Overall structure of the AlphaFold predicted VgrS:Hcp complex (left) and the VgrS crystal structure docked with Hcp (right). Inset regions highlight several conformational changes between the VgrS experimental structure and the predicted VgrS:Hcp complex at the PAAR binding site and at the domain swap.

When comparing the crystal structure aligned to the VgrS:Hcp2 AlphaFold model, the domain swap opens the base by over ∼10 Å disrupting several predicted contacts between VgrS and the Hcp (Figure 5B). This observation strongly suggests that VgrS must change conformation from open to closed to bind Hcp. Importantly, AlphaFold could not predict the exact conformation of the loops at the VgrS:Hcp interface (pLDDT <70) and only reliably modeled VgrS:Hcp2 (Figure S4C). Previous analysis suggested that the subtle sequence differences between Hcp1/2 are due to selective recruitment of Salmonella T6SS effectors associating within the lumen of Hcp1 but not Hcp2 ^2,48^, indicating that the AlphaFold predictions are not related to VgrS:Hcp binding preference.

Additionally, the VgrS:Hcp2 complex is modelled so that the PAAR-binding site at the top of the spike is reminiscent of the closed conformation in observed in VgrG1 (Figure 3A and Figure 5B) ^20^. However, the predicted complex exhibited a high predicted aligned error (PAE) (>30Å) in the loop regions of both proteins. This was not unexpected, as AlphaFold often exhibits high error for protein dynamics ^49,50^ (Figure S4C). Interestingly, regions in VgrS distant from the VgrS:Hcp interface also showed high pLDDT scores. This was unexpected as when modelled alone the VgrS β-helix neck regions is of high confidence (Figure S3B). Importantly, the unique crystal packing of VgrS is such that the PAAR binding sites of each VgrS pack directly against a PAAR binding site symmetry mate (Figure S2A). This possibly forces the PAAR binding site of VgrS to adopt the flat “PAAR-bound” conformation (Figure 3A and Figure 5B). Overall, the structural data compared to both known VgrG structures and AlphaFold predictions supports the hypothesis that VgrS changes conformation upon either PAAR or Hcp binding from open to closed.

To test that VgrS changes conformation upon binding, a structure of VgrS bound to PAAR and/or an Hcp would be required. To date we have not been able to obtain a structure of VgrS bound to its cognate PAAR domain either by X-ray crystallography or cryoEM. This is in part due to the complexity of the system as the PAAR is split and requires an Eag chaperone for both solubility and interaction with VgrS ^23^. To experimentally test Hcp binding, we were able to obtain and purify Hcp2 and assayed VgrS:Hcp2 complex formation by size-exclusion chromatography (SEC). Our binding assay was unable to show complex formation between VgrS and Hcp2 (Figure S5). This could be due to several factors, including that the PAAR domain binds VgrS first and induces the closed VgrS conformation for Hcp binding, or that the Hcp must hold an effector its lumen to bind VgrS ^25,51,52^. Alternatively, the construct cloning artifacts such as affinity-tag placement may interfere with complex formation or VgrS has higher affinity for other Hcp proteins. Regardless, the experimental structure of VgrS clearly displays a unique open conformation that based on known VgrG structural data is likely incompatible with Hcp binding.

## DISCUSSION

Understanding the T6SS is crucial for deciphering bacterial interactions with other species, especially in enteric bacteria known to cause severe disease. Here, we solve a structure of VgrS from *S. enterica* serovar Typhimurium. VgrS is a core T6SS component that serves as a modular platform for effector loading. Overall, the structure of VgrS diverges from other known VgrG proteins revealing molecular surfaces likely important for specific effector loading (Figure 1 and Figure 4). Additionally, VgrS contains a domain swap in the gp27 head domain which results in a large open conformation not previously observed in any other VgrG protein (Figure 3). Moreover, our structural data compared to previous VgrG studies indicates that VgrS may cycle between a closed conformation similar to known VgrG proteins and the conformation shown in our crystal structure (Figure 4 and Figure 5). This conformational change could be linked to a PAAR domain binding at the top of the gp5 spike, which in turn could influence the base of the gp27 head to allow for Hcp binding, or vice versa.

The VgrS crystal structure reveals an “open” conformation exhibited by the wide diameter of the gp27 cavity compared to other known VgrGs ^17,18,20^. This open conformation features a flat spike which would facilitate PAAR binding. This is opposite to the uneven surface seen in the open conformation of VgrG1, which is thought to inhibit PAAR binding. This difference in conformation suggests that the VgrS open conformation spike flat surface might allow for PAAR attachment. However, this conformation may also be due to crystal packing. As shown in Figure S2A, VgrS packs tip-to-tip at the PAAR binding surface which may influence its conformation.

The most prominent unique structural feature of VgrS is the domain swap within the gp27 head. This region is an ordered antiparallel β-sheet in all other known VgrG proteins (Figure 3). The domain swap between each monomer contributes to the open conformation observed in the crystal structure. Subsequently, the crystallographic B-factors (Figure S3D), AlphaFold modelling (Figure 5, Figure S3, and Figure S4) and a known PAAR-VgrG-Hcp complex^18^ suggest that VgrS undergoes a significant conformational change upon effector (PAAR) or Hcp binding. Namely, that the domain-swap region might relay a conformational change across the VgrS structure in response to effector loading. Given this, VgrS likely starts in the open conformation similar to our crystal structure and is unable to bind Hcp (Figure 5B). Upon loading of a PAAR or other effector at or near the C-terminal region PAAR binding site, VgrS changes to the closed conformation observed in other VgrS proteins. At this point, the gp27 base compresses and binds to either Hcp1 or Hcp2 that may already contain an effector within its ring. The VgrS:Hcp:effector complex can then nucleate assembly of the phage tail in the T6SS baseplate. Alternatively, an effector-loaded Hcp is able to bind VgrS and induce the closed conformation. These models are supported by the observation that the binding of VgrG proteins to their cognate Hcp appears to act as a checkpoint, ensuring that effectors are loaded before the subsequent polymerization of Hcp/TssBC sheath proteins and secretion out of the cell ^25^. However, our model is speculative and requires additional experimental structural data to test and observe any VgrS conformational changes.

Previous findings suggest that multiple effectors can be secreted in a single T6SS firing event, indicating that VgrGs can be decorated with multiple effectors. However, the process of loading these effectors onto VgrG is still not fully understood. The electrostatic surface of VgrS reveals distinct patches of electronegative residues on both the external and internal sides of the gp27 domain (Figure 4). The external face, known to interact with the trafficking domains of effectors like the Rhs cage, is characterized by similar electrostatic interactions ^30^. Additionally, our structural analysis of VgrS also suggests that effector loading may occur within the cavity of the gp27 head domain. Positioned at the bottom center of the trimer, this cavity forms a small triangular aperture with electrostatic residues that are accessible to the surrounding solvent (Figure 4B). This internal surface could be an effector loading site as the expansive ∼52Å wide cavity is large enough to accommodate a globular protein. Indeed, previous Cyro-EM structures of VgrGs have shown potential cargo shuttled into the cavity of the gp27 domain ^21^.

Finally, the unique C-terminal tail of VgrS is also a likely site of *Salmonella* specific effector loading. This is supported by work showing that adaptors specifically load effectors onto the VgrG tail ^22,30,42^. However, the adaptors required for this process have yet to be identified in *Salmonella.* Overall, the structure of VgrS reveals several potential sites of effector loading specific to *Salmonella* and provides a structural model of how effector loading and Hcp binding may be regulated.

## METHODS

### Construction of expression constructs

VgrS and Hcp2 from *S. enterica serovar* Typhimurium SL1344_0284, codon optimized for *E. coli* was synthesized by Genscript in the vector pET22b. VgrS was placed between restriction sites NdeI and Xho1 with a N-terminal HRV-3C 6His-Tag for purification. A truncated version of this wild-type VgrS lacking the first 81 amino acids was created by Q5-mutagenesis (NewEngland Biolabs, NEB) to generate VgrS (residues 81-729) using primer pairs 5’-CAAGATGCTGTGAGCGCGATTGTG and 5’-ATCGTCCGCGGTAACCGC. Primers were generated from Integrated DNA Technologies (IDT) and the construct was confirmed by sequencing (SickKids Toronto).The PLP Pro construct used as a positive control in our inductively-coupled mass-spectroscopy (IP-MS) experiment was borrowed from previous studies ^45^. The SciW construct used as a negative control in our ICP-MS was also from previous studies ^23^.

### Protein expression and purification

Once the expression construct was generated, the vector was transformed into *E. coli* BL21 (DE3) Gold cells which were grown in 50 mL Lysogeny Broth (LB) supplemented with 100 ug / mL ampicillin at 37 °C overnight. The following morning, overnight cultures were transferred to 3L of fresh LB broth at a 1 and 100 dilution and grown at 37 °C until an Optical Density (OD) of 0.6 at 600nm absorbance was reached. Protein expression was induced by the addition of 1 mM isopropyl β-d-1-thiogalactopyranoside (IPTG) and cells were further incubated for ∼16 hours overnight at 20 °C. After incubation, the cells were harvested by centrifugation at 4200 rpm for 30min and resuspended in 50 mL lysis buffer (50 mM Tris pH 8, 500 mM NaCl, 20 mM Imidazole). Before lysis, 1 mM PMSF and 10 mM MgCl2 were added to resuspended cells, along with a small amount of DNase. The cells were then lysed using an Emulsiflex-C3 High Pressure Homogenizer (Avestin) and centrifuged at 17,000 rpm (30,000xg) for 30 minutes at 4 °C. The supernatant was run through a nickel-NTA agarose affinity chromatography gravity column (Goldbio) equilibrated by lysis buffer. The bound protein was washed with 5x column volumes of lysis buffer before being eluted by 1x column volumes of elution buffer (50 mM Tris pH 8, 500 mM NaCl, 500 mM imidazole). The collected elution was concentrated to 2 mL by centrifugation using a 30 kDa concentrator (Amnicon) and applied to a HiLoad 16/600 Superdex 200 column (GE Health Care) for SEC using a AKTA Pure (Cytivia). The column was equilibrated with gel filtration buffer 50 mM Tris pH 8, 250 mM NaCl, 1mM 2-Beta-mercaptoethanol (BME). Hcp2 was purified as described previously, except that purification was performed in lysis buffer containing 25 mM Tris (pH 8.0), 150 mM NaCl, and 5% glycerol, followed by SEC in buffer containing 50 mM Tris (pH 8.0) and 150 mM NaCl. Hcp2 SEC buffer was also used for the SEC binding assay. To confirm purity of all protein variants, fractions were run on a 15% SDS-PAGE gel and visualized with Coomassie Brilliant Blue. Selected fractions were concentrated and used for further experiments or flash frozen in liquid nitrogen for future use.

### Size exclusion chromatography binding assay

VgrS and Hcp2 were purified individually as described above. To assess binding, the two proteins were mixed at a 1:2 molar ratio (VgrS:Hcp2) and incubated on ice for 15 minutes. The individual proteins and the mixture were then analyzed by size-exclusion chromatography (SEC) on a Superdex 200 Increase column (GE Healthcare) using an ÄKTA Pure system (Cytiva). The buffer used was the Hcp2 SEC buffer, 50 mM Tris (pH 8.0) and 150 mM NaCl. SEC fractions were also run on a 15% SDS-PAGE gel and visualized with Coomassie Brilliant Blue.

### Protein crystallization of VgrS

Crystallization of purified VgrS (81-729) with the 6His-tag was screened using commercially available screens (NeXtal, Molecular Dimensions) and an in-house Crystal Gryphon robot (Art Robbins Instruments). Conditions were screened at a protein concentration of 32 mg/mL, 18 mg/mL, and 8 mg/mL. VgrS crystals with the highest resolution of diffraction were obtained from 0.1M sodium acetate pH 5.0, 1.5 M ammonium sulfate in a 1:1 drop ratio using the sitting drop vapour diffusion method and incubated at 4 °C.

### Data collection and refinement

A VgrS data set was collected at the Canadian Light Source beamline CMCF-BM (08B1). VgrS crystals were cryo-protected by 25% glycerol. The reflections were integrated and scaled using XDS and merged using AIMLESS from the CCP4 program suite ^53,54^. Data quality was analyzed with phenix.xtriage from the PHENIX package ^55^. The structure of VgrG1 from *Pseudomonas aeruginosa* (PDB: 6H3L) was used to obtain the phases by molecular replacement using Phaser ^56^. The initial model was then refined using phenix.refine from the PHENIX package, alternating with manual building using Coot ^57,58^ The stages of refinement were done using refmacat on the CCP4 cloud ^59^. Note that the data set can be processed in P6_3_22 with a VgrS monomer as the asymmetric unit. The space group C222_1_ was chosen to have the full biological unit of a VgrS trimer in the asymmetric unit. This choice was made for biochemical relevance as VgrS has domain swaps, differences between each monomer, and has a ligand coordination site constructed in part from each monomer. Model statistics are listed in Table 1. Molecular visualizations and graphics were generated using ChimeraX.^60^

### Inductively coupled plasma mass-spectrometry (ICP-MS)

Purified SciW, PLpro, and VgrS were dialyzed overnight at 4°C into 10 mm Tris pH 7.5, 50 mM NaCl. 100 μL of each protein was added separately into 1 mL of mass-spectrometry grade nitric acid (Sigma) and 0.5 mL H_2_O_2_ (Sigma) for overnight digestion at room temperature. After digestion, milli Q water was used to dilute the digested solution to 10 mL. The same procedure was also done with 100 μL of dialysis buffer. The digested solutions were further diluted 10x for ICP-MS analysis. NIST 1643 was used as QA/QC (quality assurance/control) and indium (In) was used as an internal standard for the ICP-MS analysis. ICP-MS was carried out using an Agilent 8900 ICP-MS.

## DATA AVAILABILITY

The X-ray structure of *Salmonella* Typhimurium str. LT2 VgrS has been deposited in the Protein Data Bank under accession code 9MLP.

## SUPPORTING INFORMATION

This article contains supporting information.

## AUTHOR CONTRIBUTIONS

K.S., M.V.S., G.P conceived the study. All authors contributed to the experimental design. M.V.S and K. L. W. expressed, purified, and crystallized proteins. K.S., M.V.S. and G.P. solved and analyzed the crystal structure. K.S. and G.P. performed structural modelling. K.S., M.V.S. and G.P. wrote the paper. All authors provided feedback on the manuscript.

## ACKNOWLEDGEMENTS

We would like to thank Wei Wang in the Department of Earth Sciences at the University of Manitoba for help with running and analysis of ICP-MS data. We thank Dr. Brian Mark for providing an expression plasmid for PLpro. We would also like to thank beamline CMCF-BM at the Canadian Light Source, which is supported by the Canada Foundation for Innovation (CFI), the Natural Sciences and Engineering Research Council (NSERC), the National Research Council (NRC), the Canadian Institutes of Health Research (CIHR), the Government of Saskatchewan.

## FUNDING AND ADDITIONAL INFORMATION

This work was supported by the Canadian Institutes of Health Research (CIHR) project grant PJT-180450 to G.P. and a Research Manitoba New Investigator operating grant to G.P. K.S. was supported through a National Science and Engineering Council (NSERC) postgraduate doctoral scholarship and M.V.S was supported by a Research Manitoba Graduate Studentship and a University of Manitoba Graduate Fellowship.

