## Supplemental_data for "Structure of the type VI secretion protein VgrS from *Salmonella* Typhimurium"

* Indicates equal author contribution

**Keywords:** Type VI Secretion System, VgrG, *Salmonella* Typhimurium, Bacterial warfare, Bacterial Secretion, X-ray Crystallography.

#To whom correspondence should be addressed: GP

Telephone – 204-474-6543

List of Supplementary information:

Figures S1-S5

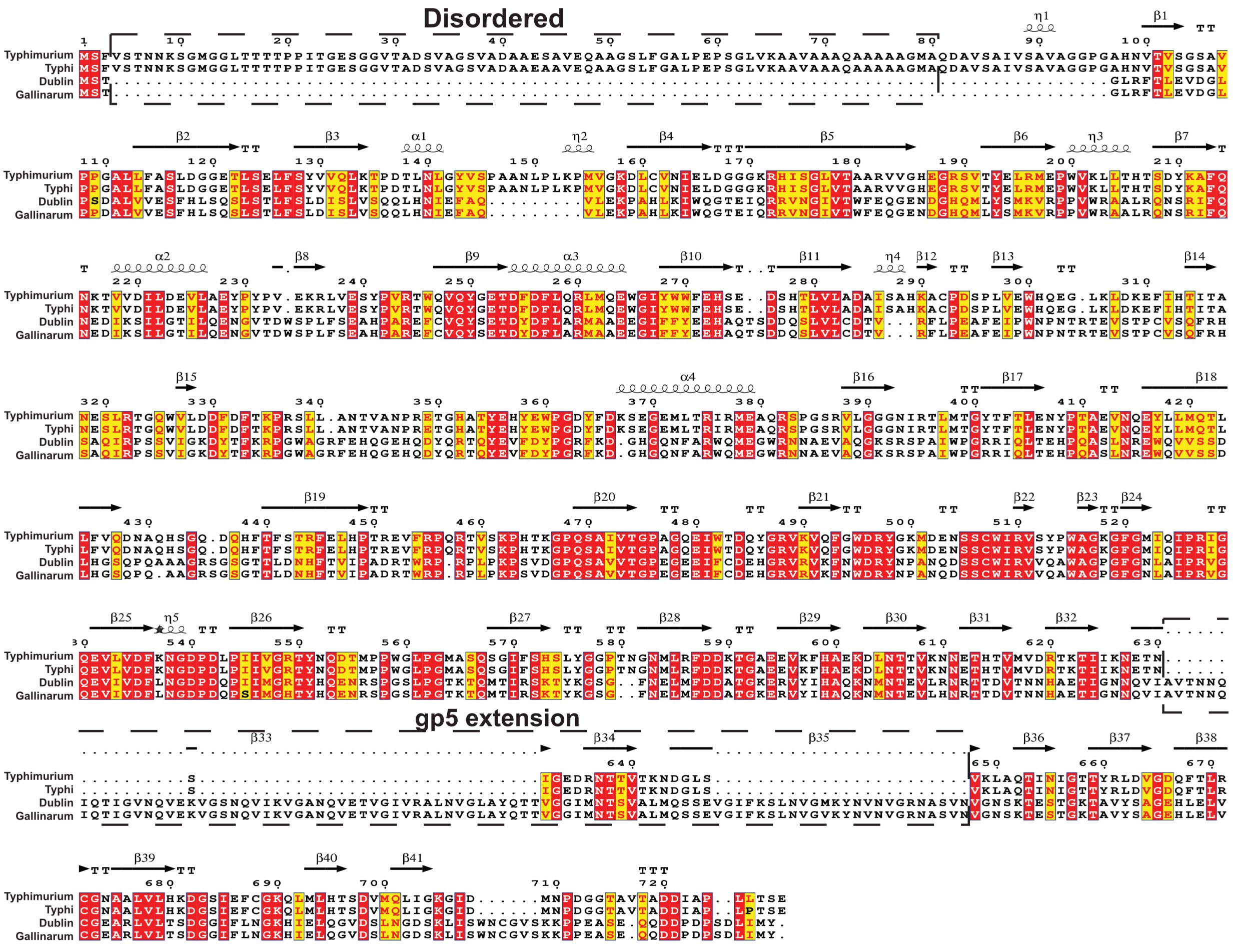


**Figure S1: Sequence alignment of the VgrS proteins from Figure 1B.** Alignment was made with clustalo and colored by Espript3.

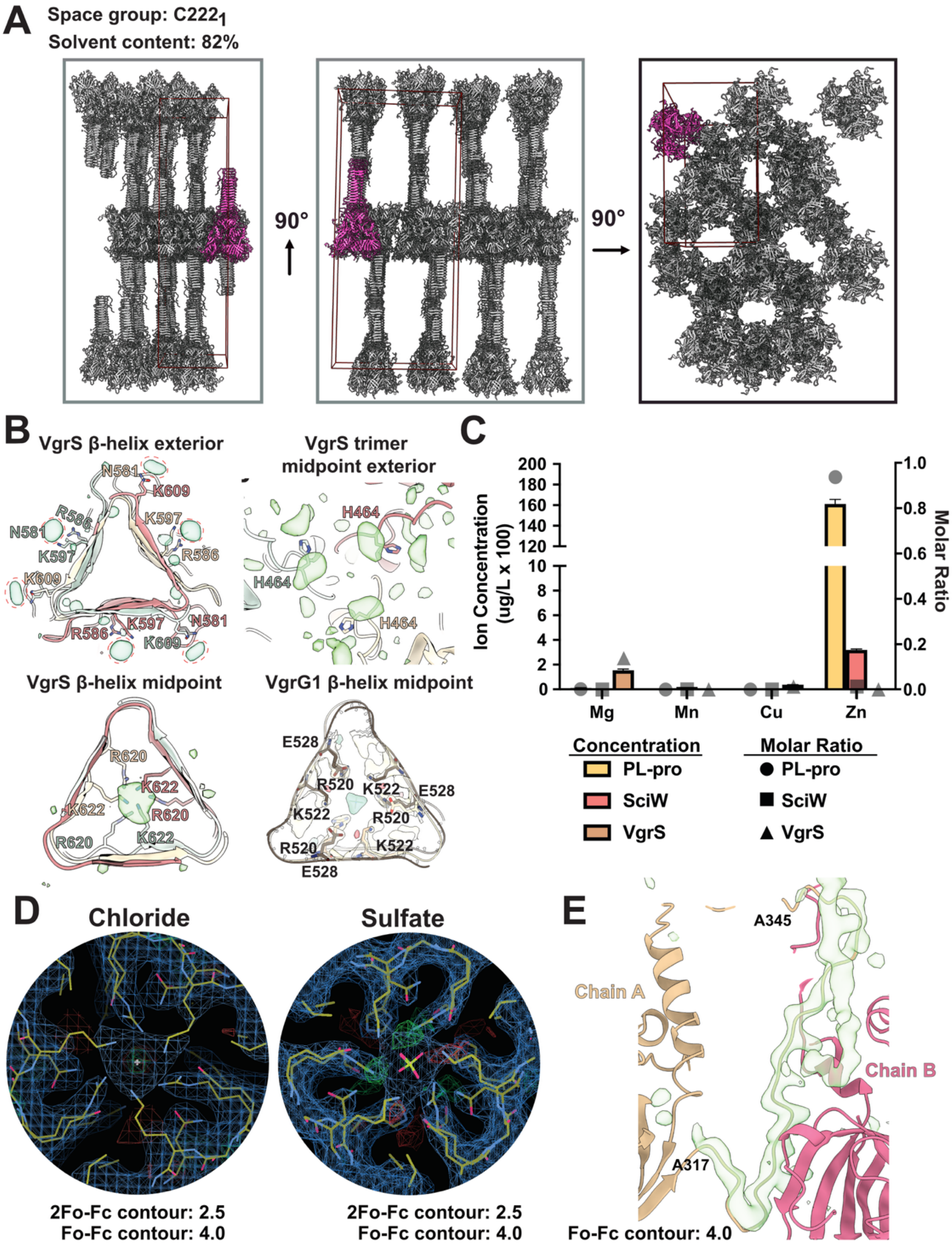


**Figure S2: Packing and electron density analysis for VgrS. A)** Crystal packing of VgrS in the C222$_1$ space group. The VgrS trimer of the asymmetric unit is shown in magenta **B)** The observed electron density from mFo-DFc maps contoured at 4.0 rmsd and drawn in green. Residues interacting with potential ligands are highlighted. VgrG1 is PDB: 4UHV. **C)** ICP-MS detection and quantification of metal ions within VgrS (brown), SciW (red) and the zinc binding protein MersPLpro (gold). Measured concentration of ions from each sample (10 mg/mL protein) is shown on the left Y-axis. The molar ratio of ion to protein concentration is shown on the right Y-axis corresponding to the symbols. VgrS (triangle), SciW (square), PLpro (circle). **D)** 2mFo-DFc and mFo-DFc map of sulfate and chloride ions being modelled into the electron density within VgrS β-helix. **E)** Calculated mFo-DFc map shown in green with the domain swap region residues A317 to A345 deleted.

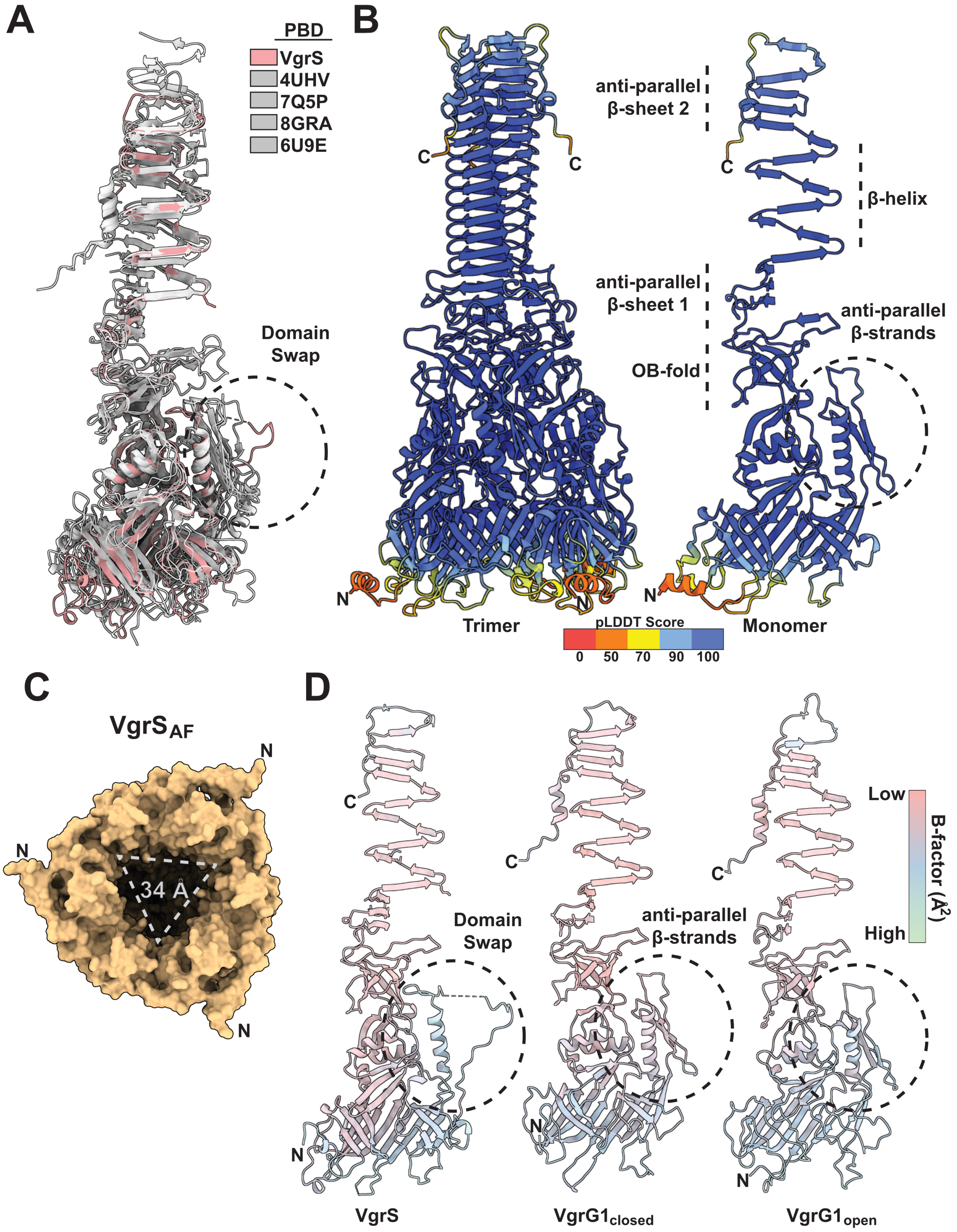


**Figure S3: Structural comparison of VgrS with other known VgrGs. A)** Overlay of the VgrS monomer with other known VgrG monomers (PDB: 4UHV, 7Q5P, 8GRA, 6U9E).

The domain swap found in VgrS is indicated by dashed circle. **B)** AlphaFold3 prediction of VgrS colored according to pLDDT error score. **C)** Bottom view of $VgrS_{AF}$ with triangular dimensions shown indicates a closed conformation similar to VgrG1 **D)** B-factors of VgrS and VgrG1 in two different conformations (PDB: 4UHV).

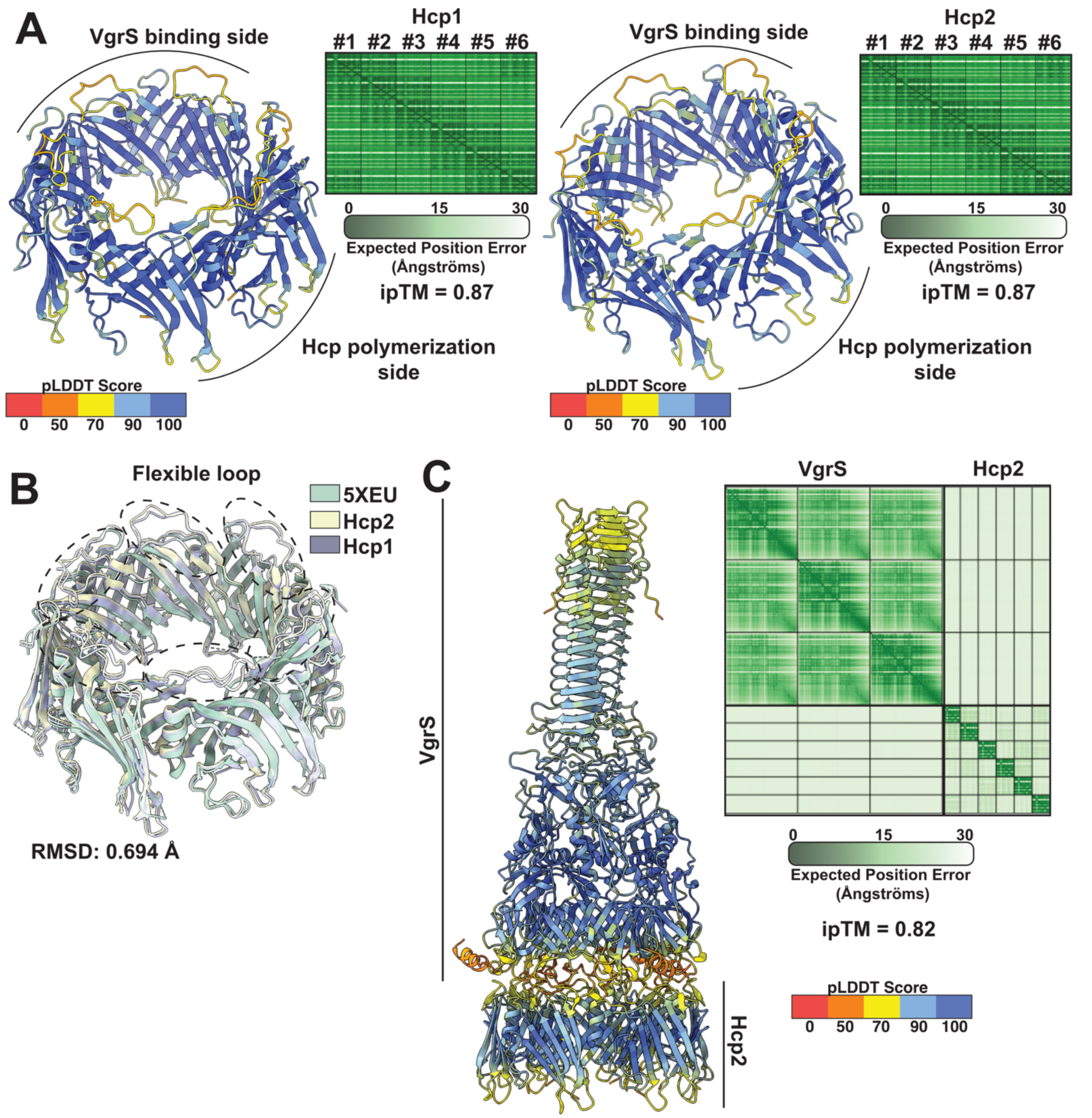


**Figure S4: AlphaFold3 predictions of *Salmonella* Hcps and a VgrS:Hcp complex. A)** AlphaFold3 predictions of hexameric Hcp1 (STM0276) and Hcp2 (STM0279) with corresponding PAE plots and iPTM scores. **B)** Overlay of the Hcp1 and Hcp2 AlphaFold predictions with the known structure of Hcp2 (5XEU). **C)** AlphaFold3 prediction of VgrS bound to Hcp2 with corresponding PAE plot on the right. The model is colored based on pLDDT error scores and highlights significant uncertainty within the gp5, domain swap, and interacting loops of VgrS and Hcp2. VgrS:Hcp2 ipTM was only 0.52 so is not shown.

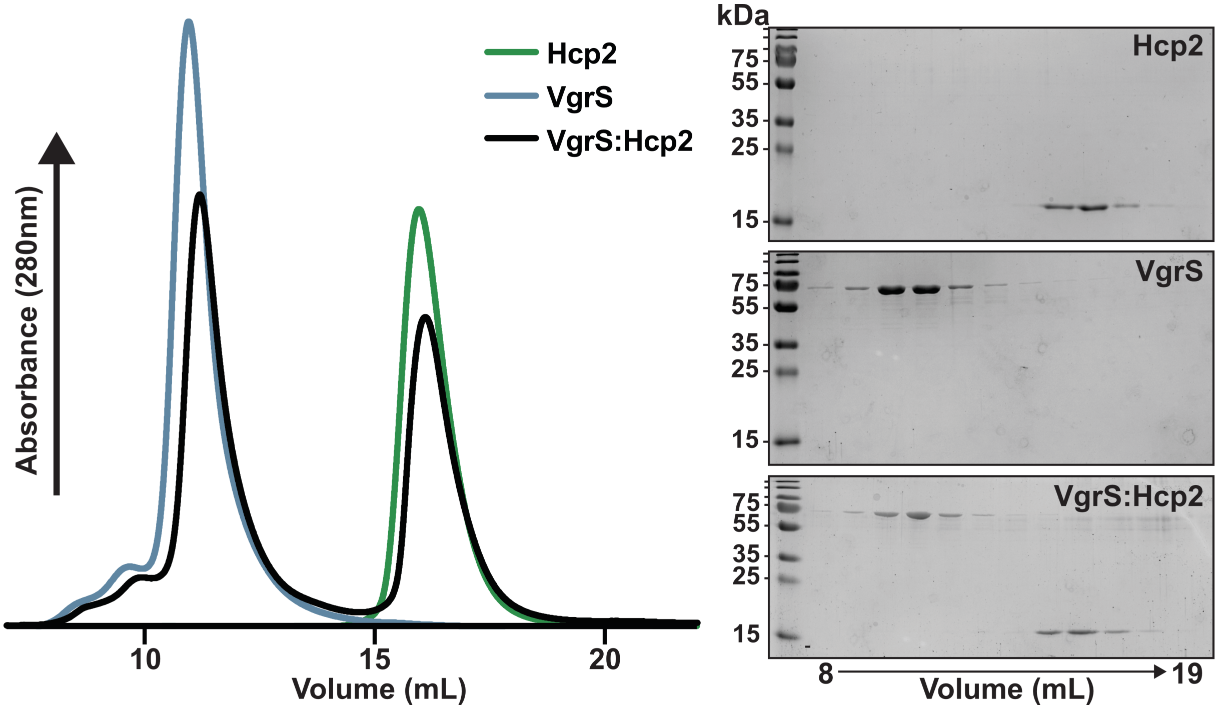


**Figure S5. Size-exclusion binding assay with purified VgrS and Hcp2.** Left: Size-exclusion chromatogram of VgrS and Hcp2 run individually or as a mixture (1:2 ratio) on a Supdex200 Increase column. Right: Coomassie stained SDS-PAGE gels of fractions collected from each SEC binding assay.
